# Gene Flow and Recent Lineage Colonization Constrain Genetic Differentiation Despite Local Adaptation

**DOI:** 10.64898/2026.08.15.745018

**Authors:** Dylan J. Padilla Perez, Lisa K. Brady, Jody M. Taft, Nathaniel B. Edelman, A. Z. Andis Arietta, David K. Skelly

## Abstract

Demographic processes such as colonization to new environments and gene flow fundamentally shape the genomic landscape, either facilitating or constraining the efficiency of selection by altering the balance between genetic diversity and adaptive responses. Although theoretical predictions suggest that the efficacy of selection is dictated by a species’ demographic history, empirical studies often overlook these constraints, yielding misleading observations. In this study, we present the first functional genome annotation for the wood frog (*Lithobates sylvaticus*), providing a critical genomic resource for understanding the adaptive capacity of the species. Based on the annotation, we examined the potential for selection to drive genomic and phenotypic divergence among populations distributed across vernal ponds in Northeastern Connecticut, USA. A genotype*×*environment association analysis revealed that the frequency of an outlier loci (*Rab28*) spikes in response to one wetland that is notable for having relatively low canopy cover and large area. We also found that selection has driven a strong disparity in embryonic development among populations of wood frog at a rate exceeding that of neutral genetic drift. This genomic signature of selection together with a remarkable phenotypic differentiation suggests that natural selection overcomes the power of genetic drift, even in a landscape characterized by relatively recent colonization and substantial evidence of connectivity among breeding wetlands. These findings improve our understanding of the wood frog’s variation at a microgeographic scale.

## Introduction

The structuring of genetic variation within and among populations is fundamentally driven by a complex interaction among evolutionary forces. While neutral forces such as genetic drift could press populations to diverge by allowing for the stochastic accumulation of allelic differences, the role of natural selection is recognized as a primary driver of genomic differentiation (Nielsen, 2005). Under this framework, natural selection increases the frequency of alleles that confer fitness advantages in specific environments, potentially leading to the formation of “genomic islands of speciation” where differentiation exceeds the background levels established by neutral forces (Wolf and Ellegren, 2017). Advances in population genomics now allow for fine-scale mapping of these signatures of selection, providing unprecedented insights into the means by which ecological pressures reshape the genetic architecture of natural populations (Savolainen et al., 2013). Understanding this process is essential for accurately assessing the capacity of populations to adapt to rapidly changing environments.

While the drift-selection balance facilitates genetic structure among populations, other evolutionary forces maintain genetic cohesion, often resulting in relatively low genetic differentiation. One countering force is the rate of gene flow among populations, which can swamp the effects of local adaptation by homogenizing allele frequencies across environments (Lenormand, 2002). When the strength of gene flow (*m*) significantly exceeds the strength of selection (*s*), adaptive alleles might fail to establish and negligible divergence is expected, especially among small populations (Richard-son et al., 2014; Lenormand, 2002). For example, studies in the European Grayling (*Thymallus thymallus*) show that strong gene flow across rivers hinders population structure even when there is rapid adaptation to local temperatures, showing that *m* can prevent long-term differentiation (Junge et al., 2011). Even when selection is sufficient to overcome the effect of genetic drift, divergence takes time. In some systems, initial colonization is recent, meaning that there may not have been sufficient time to allow divergence, thereby preserving substantial ancestral polymorphism (Mila et al., 2007). For instance, in species that have undergone a series of rapid colonizations such as the Silvereye (*Zosterops lateralis*), populations exhibit negligible differentiation at neutral loci as a result of the extremely short timeframe for genetic drift to operate (Clegg et al., 2002). However, the lack of substantial neutral differentiation does not necessarily imply a lack of adaptive divergence among phenotypic traits that are critical for survival and the persistence of species (Nosil et al., 2009).

A pronounced phenotypic divergence in the absence of genome-wide genetic differentiation might be observed in species that undergo rapid ecological adaptation. This phenomenon often arises when strong disruptive selection operates on specific quantitative traits (Merilä and Crnokrak, 2001). To estimate the disparity between phenotypic divergence and genetic divergence, researchers compare the degree of quantitative trait differentiation (*Q_st_*) with the differentiation at neutral molecular markers (*F_st_*). Under neutral evolution, *Q_st_* is expected to equal *F_st_*. However, when *Q_st_ > F_st_*, it provides robust evidence that divergent selection is driving phenotypic local adaptation at a rate exceeding that of neutral genetic drift (Whitlock, 2008). A compelling example is found in the common frog (*Rana temporaria*), where populations across steep latitudinal gradients exhibit marked differences in metamorphic traits, such as growth rate and development time, to cope with shorter growing seasons. Despite these significant life-history shifts (*Q_st_*), neutral genetic markers (*F_st_*) show negligible differentiation, indicating that selection for developmental timing has rapidly outpaced the effects of genetic drift in these recently established post-glacial populations (Palo et al., 2003). Similarly, studies on the Threespine stickleback (*Gasterosteus aculeatus*) demonstrate that while marine and freshwater populations show minimal differentiation at neutral loci due to recent colonization, they exhibit high *Q_st_* values for morphological traits like armor plating, reflecting intense selection for different ecological niches (Leinonen et al., 2006).

The wood frog (*Lithobates sylvaticus*) presents a particularly compelling system to examine forces that shape genomic and phenotypic evolution among populations. In Northeastern Connecticut, USA, the Yale-Myers Forest offers us the opportunity to study populations of wood frogs that breed within vernal wetlands (also referred to as “ponds”) that are known to dry on an annual basis, and to vary drastically in environmental conditions (Figure 1). For example, owing to woody vegetation dominated by Red Maple (*Acer rubrum*) and Eastern Hemlock (*Tsuga canadensis*), the canopy cover percentage among wetlands can range from 10% to 89% (Skelly and Freidenburg, 2000). Because wood frogs are known for having high breeding-site fidelity, particularly once an individual has reached sexual maturity, we assumed that each wetland was occupied by a unique population (Berven and Grudzien, 1990). When experimentally exposed to canopy cover conditions that mimic those experienced in the forest, larvae of the wood frog that occupy closed-canopy wetlands select warmer temperatures than those expected by chance (Kealoha Freidenburg and Skelly, 2004). By contrast, embryos in the closed-canopy wetlands took up to twice as long to hatch compared to those that occupy opened-canopy wetlands (Skelly, 2004), and that pattern of microgeographic variation still persists in the same populations nearly two decades later (Arietta and Skelly, 2021). Although these studies have improved our understanding of the factors that lead to drastic phenotypic variation, we know little about the factors that potentially cause genetic variation among populations of the wood frog. Here, we detect selection signatures across the genome of the wood frog, and estimate some demographic processes, such as gene flow and recent colonization, to provide a robust framework that enables us to elucidate signals of genomic and phenotypic evolution. To do that, we provide a detailed genome annotation of the wood frog, establishing a robust genomic tool necessary to distinguish signatures of selection from noise in a relatively large, complex amphibian genome. By employing the annotation as a diagnostic tool and reconstructing the demographic history of the species, we identify signatures of selection across populations, testing the hypothesis that phenotypic variation does not necessarily reflect strong genetic structure.

**Figure 1:**
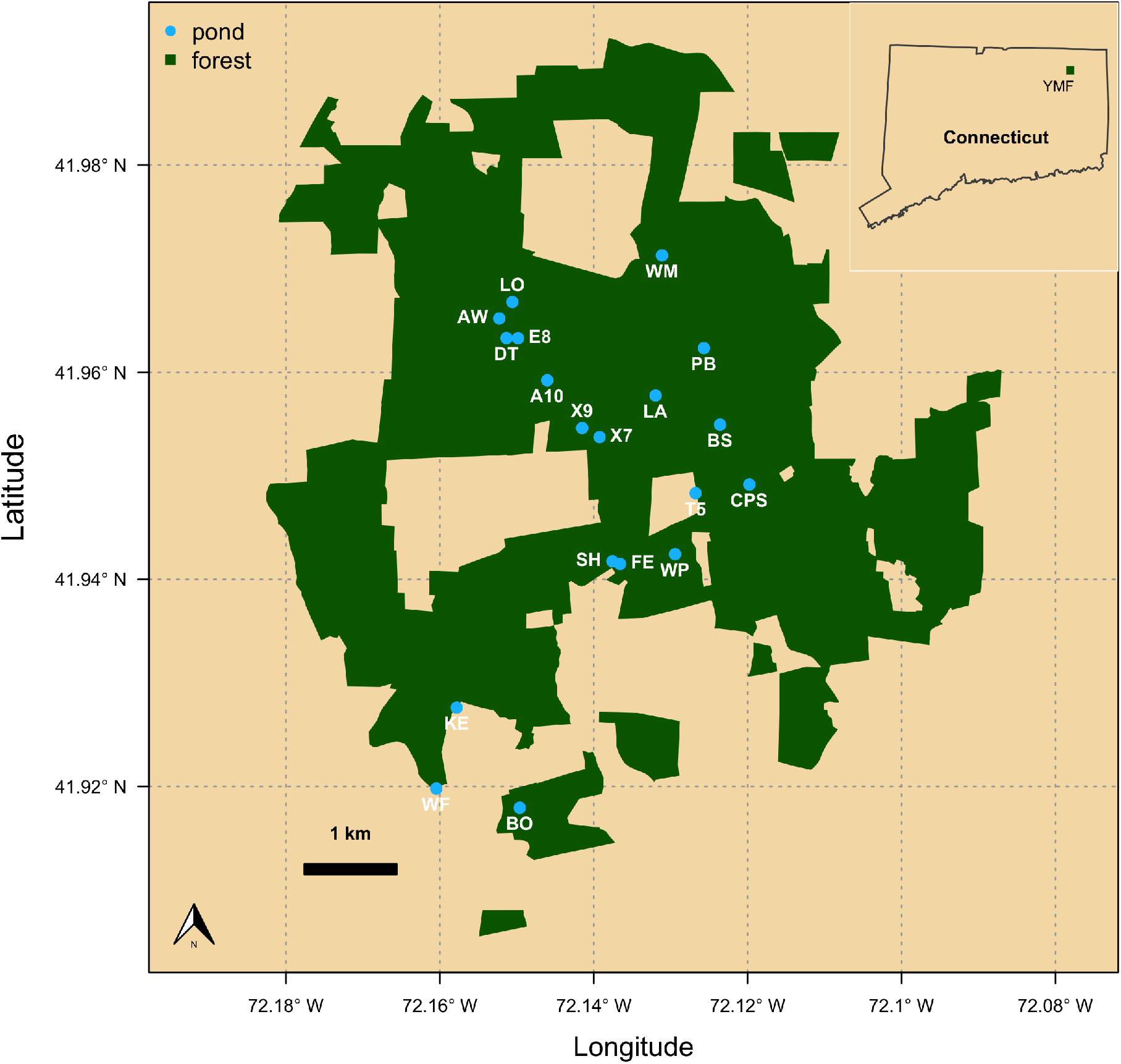
Geographic distribution of study wetlands (populations) within the Yale Myers Forest, Connecticut, USA.

## Materials and Methods

### Wood frog genome annotation

We employed the genome annotation based on the current wood frog genome assembly available on NCBI (GCA_028564925.1_aRanSyl1). We used the GALBA v1.0.11 pipeline for de novo gene predictions with AUGUSTUS and Miniprot (Bruuna et al., 2023; Li, 2023; Buchfink et al., 2015; Stanke et al., 2006; Hoff and Stanke, 2019). To meet the input requirements of the GALBA pipeline, we used Repeat Masker 4.1.2.p1 https://www.repeatmasker.org/, which enabled us to soft-mask all repetitive elements in the assembly. We also used protein sequence data from the American Bullfrog (*Aquarana catesbeiana*: GCA_042186555.1), the Hourglass Treefrog (*Dendropsophus ebraccatus*: GCF_027789765.1), and the Common Frog (*Rana temporaria*: GCA_905171775.1) as protein sequence evidence data in the training of AUGUSTUS. We assessed the completeness of the annotation using BUSCO in protein mode https://busco.ezlab.org/. For functional annotations of the predicted proteins, we used BlastP to search against the NCBI-BLAST protein database (Camacho et al., 2009) with an e-value cutoff of 10*^−^*^4^ and *−max*_*target*_*seqs* argument set to 10. We used Annie v1.0 (Tate et al., 2014) to integrate functional evidence from SwissProt BLAST results, assigning standardized gene names and product descriptions to predicted features. ANNIE produced a three-column table linking feature IDs to their corresponding annotations for downstream integration with the General Feature Format file (GFF3). We merged functional gene annotations produced by the ANNIE pipeline with the structural annotation in GFF3 format using a custom Python script. The script parses the ANNIE output (three-column TSV containing feature identifiers, annotation field names, and values) and systematically integrates the corresponding information into the GFF3 attribute field. Specifically, the script associates each feature’s ID in the GFF3 file with matching name and product entries from the ANNIE output. Where available, product descriptions were appended to both gene and mRNA features, and a name tag was added or updated to reflect the human-readable gene symbol. All attribute values were percent-encoded to ensure GFF3 format compliance. Parent-child relationships and all original identifiers were preserved, allowing direct correspondence between the structural and functional annotations. We then filtered the gene models in a stepwise process, first retaining all gene models with orthologs, using Protein-Protein BLAST 2.12.0 (Camacho et al., 2009), and then manually filtering all remaining unnamed gene models (those without a BLAST hit), only including those with at least two exons in the final annotation. Final annotation statistics were generated using AGAT v1.4.2 (Dainat et al., 2020) and BUSCO.

### Tissue sampling, DNA extraction, library preparation, and sequencing

In 2018, we assembled a comprehensive dataset of DNA samples collected from wood frog populations distributed across 19 wetlands at Yale-Myers forest (Figure 1). The samples were preserved in 90% Ethanol and brought to the laboratory, where we isolated genomic DNA by using the DNeasy Blood and Tissue Kit (Qiagen, Germany) according to the manufacturer’s instructions. Specifically, we pre-heated the AE elution buffer to 56*^◦^*C, extended the incubation period to 10 minutes, and performed two sequential elutions using 100 *µ*L of buffer rather than a single 200 *µ*L volume. Furthermore, we consolidated two independent extractions during the final elution stage to maximize the total DNA concentration. We quantified the resulting DNA using 2 *µ*L of template on a Qubit 4 Fluorometer (Thermo Fisher Scientific, USA) and stored the extracts at *−*80*^◦^*C until processing. Samples meeting a minimum concentration threshold of 25 ng/*µ*L were transported on dry ice to the University of Minnesota Genomics Center for library preparation. We prepared double-digest RAD (ddRAD) libraries for 265 samples using a BamHI + NsiI enzyme combination, then pooled and sequenced these across two lanes of a NovaSeq SP platform (1 *×* 100 *− bp*). While we targeted a depth of approximately 2.75M reads per sample, the final sequencing yielded a mean of *≈* 3*M* reads per library with a minimum quality score of Q30 for all samples. We adhered to animal handling and tissue collection guidelines approved under IACUC protocol 2019-10361.

### Bioinformatic analyses

We processed the raw reads using the ipyrad v.0.9.107 pipeline (Eaton and Overcast, 2020). Reads were then filtered to remove adapters and low-quality data, excluding any reads with more than five bases having a Phred quality score below 20. We also enforced a minimum post-trimming length of 35 bp. To ensure the efficacy of these steps, we double-checked the trimmed data based on a MultiQC report (Ewels et al., 2016), and subsequently filtered out any samples with high levels of adapter content. Using the reference-based assembly method, we aligned the cleaned reads to the wood frog reference genome (GCA_028564925.1_aRanSyl1). We called within-sample consensus sequences using a minimum depth of 6 reads for both statistical and majority-rule base calling, while excluding clusters with a depth exceeding 10,000 reads to avoid potential paralogs. To ensure data quality, we filtered consensus sequences to allow a maximum of 5% uncalled or ambiguous bases (Ns) and 5% heterozygous sites. Finally, we retained loci only if they occurred in at least four samples and met our thresholds for genomic variation: a maximum of 20% SNPs, 5 indels per locus, and 50% shared heterozygosity. The pipeline generated several output formats, including Phylip, Structure, and loci files; however, we primarily used the VCF (Variant Call Format) file for all subsequent downstream analyses.

To prevent skewed results caused by missing data, we filtered out any sample or SNP loci exhibiting a missing genotype rate exceeding 20%. To do so, we used the missingno function within the R package poppr v2.9.3 (Kamvar et al., 2014). Subsequently, we evaluated SNP markers for departures from Hardy-Weinberg equilibrium (HWE). We used the function hw.test from the pegas package v1.1 (Paradis, 2010), and ran 1,000 Monte Carlo permutations to generate p-value estimates. To determine which loci were out of equilibrium, we applied a False Discovery Rate (FDR) correction to the p-values; loci were subsequently excluded if they showed significant deviations in over half of the sampled populations.

### Population structure and genetic difierentiation analyses

To understand how individuals are genetically related, we examined their genetic structure and differentiation. To infer genetic structure, we estimated admixture proportions across our samples. To do this, we employed the sparse non-negative matrix factorization algorithm (sNMF) via the snmf function in the R package LEA v3.2.0 (Frichot and François, 2015). Much like Bayesian clustering approaches, this method derives individual admixture coefficients from an allele frequency matrix. The sNMF algorithm produces least-squares estimates of ancestry proportions for a predefined number of ancestral populations (*k*). To estimate the optimal *k* value, we evaluated the model’s predictive accuracy using a cross-validation-based entropy criterion. We tested a range of ancestral populations from *k* = 1 to *k* = 10, executing 50 independent iterations for each value of *k* to ensure stability and calculate mean cross-entropy.

To estimate genetic differentiation among populations, we calculated pairwise *F_st_*estimates following the Weir and Cockerham (1984) framework. This metric partitions genetic variance into within-population and among-population components (Wright, 1931). Generally, high migration rates between large populations result in negligible differentiation (*F_st_ ≈* 0), whereas restricted gene flow between small populations leads to high differentiation (*F_st_ ≈* 1). We implemented these calculations using the gl.fst.pop function within the R package dartR v2.7.2 (Gruber et al., 2018). To assess statistical significance, we performed 1,000 bootstrap replicates across all loci to derive 95% confidence intervals. *F_st_* values were categorized as significantly different from zero only if their respective confidence intervals excluded zero.

### Demographic history inference

We inferred the demographic history of the wood frog by employing a composite likelihood approach using fastsimcoal2 (Excoffier et al., 2021). Our analysis focused on five randomly selected populations (BS, DT, E8, PB, and X7) from the initial 20 populations involved in the study. We did so to maintain a balance between model complexity and computational performance, which enabled us to avoid poor parameter estimates. fastsimcoal2 uses a multidimensional Site Frequency Spectrum (SFS) as input data, which represents the distribution of allele frequencies across populations and serves as raw material for demographic inference. Since different historical events, such as bottlenecks, expansions, or migration, leave distinct signatures on the SFS, fastsimcoal2 can estimate parameters by maximizing the probability of observing the empirical SFS under a given model (Excoffier et al., 2021). To generate a multidimensional SFS from our data, we first minimized the effects associated with physical linkage across SNPs. Accordingly, we performed a linkage disequilibrium (LD) pruning on the data using PLINK v1.9.0-b.8 (Purcell et al., 2007). In doing so, we examined loci across a sliding window approach (window size: 50 SNPs; step size: 5 SNPs) and removed any SNP with *r*^2^ *>* 0.5 with any other SNP within the window. After we filtered the data to retain independent SNPs, we used easySFS (Gutenkunst et al., 2009; Coffman et al., 2016) to find the most effective population size projections for constructing the site frequency spectrum and subsequently run fastsimcoal2.

We ran fastsimcoal2 to fit and compare two competing demographic scenarios: (1) recent split with gene flow, and (2) recent split without gene flow. The parameters estimated in the models included historical effective population size (*N_e_*), migration rate (*m*), and time of divergence (*T_div_*). To convert these estimates into absolute units, we adopted a mutation rate of 1.3 *×* 10*^−^*^9^ per site per generation. We adopted this value based on representative mutation rates for other amphibian and ranid species (e.g., Crawford, 2003; Sun et al., 2015), which generally suggest slower substitution rates in amphibians compared to other vertebrates. For each model, we performed 50 independent runs with 1,000 coalescent simulations to ensure convergence. We compared the fit of the models based on *AIC* values and all downstream inferences were based on the most likely one.

Based on the most likely demographic model, we estimated the time required for two randomly selected populations (DT and E8) to reach a specific genetic differentiation (*F_st_*). Such analysis enabled us to understand how gene flow modulates the genetic structure of wood frogs’ populations. The analysis consisted of monitoring pairwise *F_st_* between the two populations over time from a simulated 10 MB genomic segment assumed to have a mutation rate of 1.3 *×* 10*^−^*9. This genomic segment meant to represent a subset of the much larger genome size of the wood frog (5.2 GB). We ran the simulation in SLiM v.5.1 (Haller et al., 2026) with the divergence time, effective population sizes, and migration rate estimated by fastsimcoal2, and a baseline recombination rate of 1 *×* 10*^−^*8.

### Detecting signatures of selection across the genome

To detect signatures of local adaptation across the genome of the wood frog, we performed a genotype*×*environment association analysis based on a latent factor mixed model (LFMM; Jumentier et al., 2022). To do this, we ran the function lfmm available in the R package LEA v.3.22.0 (Frichot and Francois, 2015). Specifically, we fitted a model informed by a matrix of allele frequencies across populations as the dependent variable, and a matrix of independent variables consisting of 4 environmental factors measured across the ponds occupied by the populations examined in this study. The environmental factors included area of the ponds, distance to closest road, elevation, and weighted global site factor (wGSF). The LFMM also required an estimate of the number of ancestral populations in the data (*k*), which relied on the admixture analysis described earlier. We identified potential SNPs under selection after controlling for Type I errors. We did so by calculating *q*-values from the observed *p*-values estimated by the LFMM. This procedure limits the expected ratio of erroneous rejections to the total number of significant findings (Storey and Tibshirani, 2003).

We used the GFF file generated from the wood frog genome annotation described above to extract gene names associated with the candidate loci under selection. We then computed the frequency of these genes across ponds to better understand their association with the environment. By comparing the overlap between gene frequencies and environmental gradients, we identified the environmental factor acting as strong selective agent.

### Quantitative trait difierentiation (Q_st_)

Finally, we compared the divergence of a phenotypic trait such as embryonic development rate, with the total genetic differentiation observed across populations of the wood frog (overall *F_st_*). This comparison enabled us to distinguish between natural selection and genetic drift as causes of population differentiation in a phenotypic trait (Leinonen et al., 2013). To do so, we used published estimates of embryonic period from populations of wood frogs collected at Yale Myers Forest within 24 hours of oviposition (Arietta and Skelly, 2021). To estimate embryonic period, the authors split clutches from all population across two temperature treatments (high:13.7°C; low: 11.7°C) and monitored embryos daily until hatching. This experimental designed enabled us to partition the total genetic variance for embryonic development into two main components: the variance among populations and the variance within populations. To do this, we modeled the fixed effect of temperature on embryonic period with the lmer function available in the R package lme4 v.1.1.38 (Bates et al., 2015). In the model, we treated population (1 | Population) and clutch nested in population (1 | Population:Family) as random effects to estimate their variance components. Subsequently, we plugged in the variance components in the *Q_st_* formula as follows:

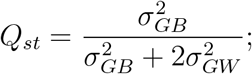

Where 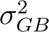 represents the genetic variance among populations and 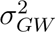 the additive genetic variance within populations.

To provide a thorough comparison, we pruned our data to only include the populations examined in Arietta and Skelly (2021). Furthermore, we ran 1000 boostrap iterations to compute 95% confidence intervals for *Q_st_* and *F_st_*.

## Results

The GALBA-predicted gene models through protein-protein BLAST ortholog retention and manual curation significantly improved the structural integrity of the wood frog genome annotation. This process reduced the initial gene count from 51,008 to 39,847, representing a 22% reduction. This contraction was primarily driven by the removal of low-evidence, single-exon predictions, which decreased from 17,940 (35% of the total) to 10,575 (27% of the total). Despite the reduction in total features, the structural quality of the remaining models improved markedly. The mean number of exons per mRNA rose by 20% (from 5.0 to 6.0), and the average gene length increased by 24% to 43,699 bp. Mean coding sequences length also grew by 14% to 1,362 bp, while total genome coverage by genes remained stable at 33.7% of the estimated 5.17 Gbp assembly. In addition, the protein-mode BUSCO analysis confirmed high annotation completeness, comprising 68.9% single-copy and 19.6% duplicated orthologs (Figure S1). The low fragmentation rate (6.3%) demonstrates that the filtering strategy successfully eliminated spurious models while preserving conserved ortholog content. Functional annotation reached a high level of coverage, with 88.53% of predicted models receiving BlastP/ANNIE assignments, including standardized gene names and product descriptions. While the current lack of RNA-seq data limits the evaluation of transcript-supported isoforms, the resulting gene set demonstrates the robustness and functional depth required for comprehensive downstream comparative genomics (see supplementary materials).

The quality control procedure of the raw reads resulted in a dataset of 15,524 loci (SNPs), and 31,048 alleles from 139 individuals. According to the snmf run from the admixture analysis, these individuals originated from a single ancestral population (*k* = 1). This analysis reveals negligible population structure and potentially high levels of gene flow across populations (Figure 2A). Pairwise *F_st_* estimates are consistent with the pattern suggested by the admixture analysis (Figure 2B), with relatively low values ranging from 0 to 0.03. Most populations remain genetically similar from each other except for a few that maintain a subtle differentiation (*F_st_* = 0.03).

**Figure 2:**
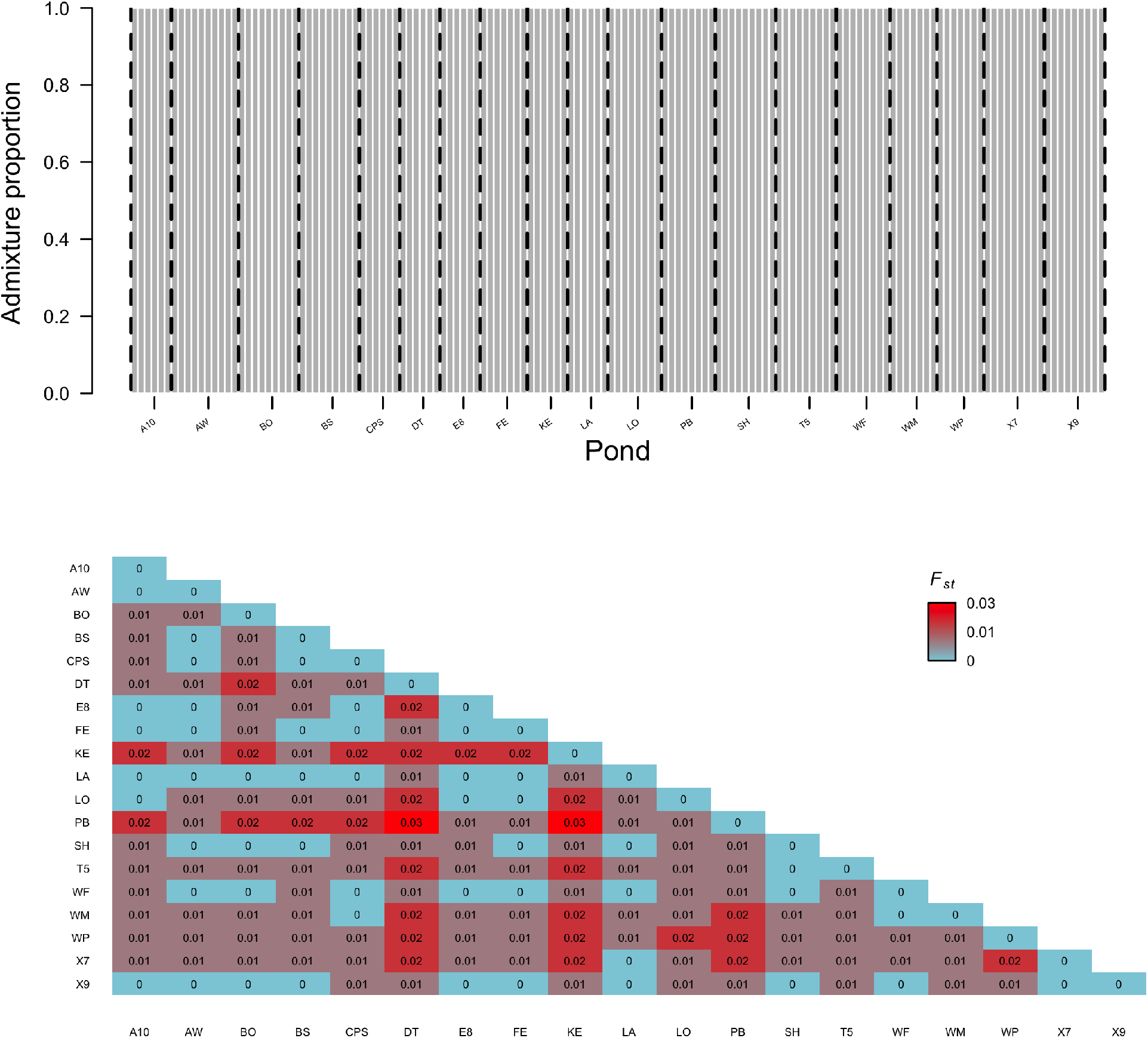
Population genetic structure and differentiation across 19 populations of the wood frog. A) Structure plot representing the admixture proportions for individuals sampled at each pond. Each vertical bar represents a single individual, with colors indicating the probability of assignment to ancestral clusters (*k*). The dashed line represents boundaries between populations. B) Heatmap of pairwise *F_st_* values, representing genetic differentiation between populations.

The genotype*×*environment association analysis reveals one outlier loci (*Rab28*) on chromosome 1 (Figure 3A). The spatial frequency distribution of this allele across populations suggests that the environmental conditions at Yale-Myers are rather heterogeneous. While the candidate allele remained at low frequencies in most ponds, we observed that its frequency spikes at pond PB, reaching a value of 0.18 (Figure 3B). These localized frequency peak at pond PB corresponds to an environmental extreme characterized by a relatively low canopy cover (wtGSF) and large area (Figure 3C).

**Figure 3:**
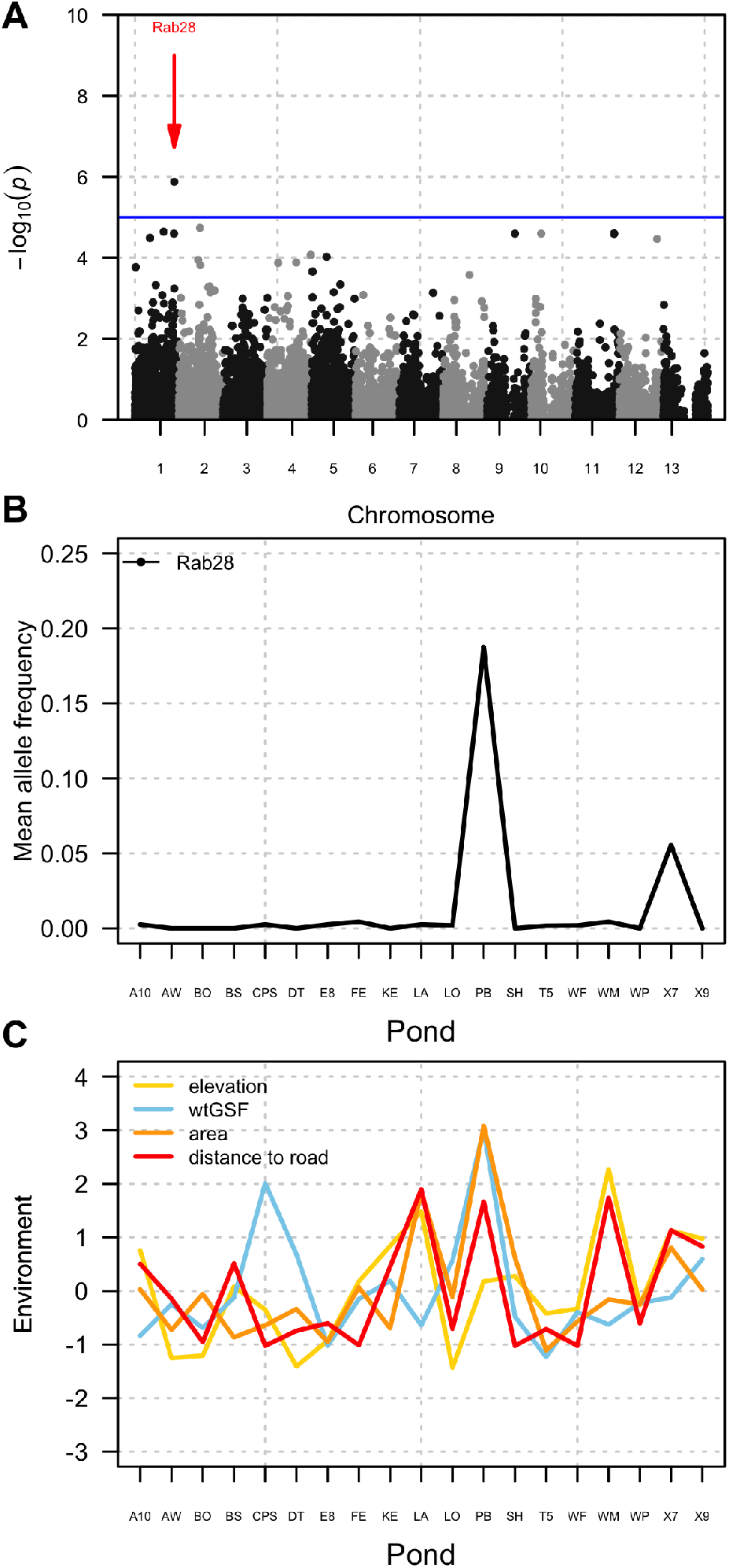
Genome*×*environment association analysis. A) Manhattan plot of genomic differentiation. The blue horizontal line represents the significance threshold for outlier detection. A red arrow highlights a significant outlier allele. B) Mean allele frequency by pond. Frequency of an outlier allele across ponds. C) Environmental variation across ponds. Standardized environmental variables across all study sites.

The demographic model suggests a recent divergence of five populations from a common ancestor approximately 24 generations ago (Figure 4A). In addition to this recent split, the model identifies a complex dynamic of contemporary gene flow, with migration rates (*m*) ranging from 4.1 *×* 10*^−^*^4^ to 1.3 *×* 10*^−^*^2^. Notably, the highest migration rates are observed from PB into DT (*m* = 1.3 *×* 10*^−^*^2^) and from BS into E8 (*m* = 4.5 *×* 10*^−^*^3^), indicating a strong genetic exchange between them. When tracking the trajectory of pairwise *F_ST_* between populations DT and E8, the SLiM simulation shows a steady increase in genetic differentiation over time, reaching a *F_st_* of 0.085 by 480–500 generations post-split. While the *F_st_* values approach the arbitrary threshold of 0.09 (dashed red line), the rate of differentiation begins to plateau after roughly 400 generations, potentially reflecting the counteracting influence of migration rates identified in the demographic model (Figure 4B).

**Figure 4:**
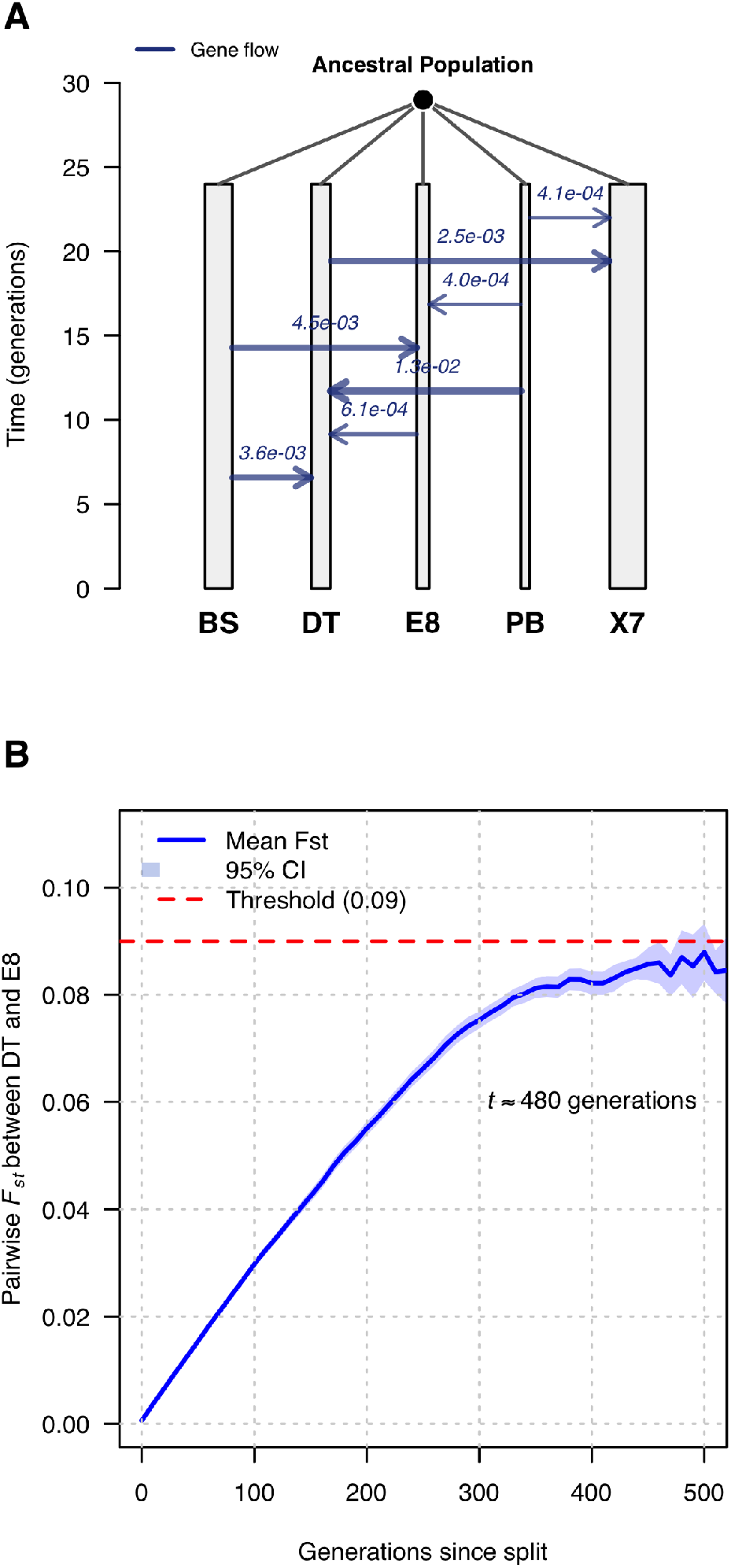
Demographic modeling and simulation of genetic differentiation over time. A) A schematic representation of the fastsimcoal2 demographic model showing the recent divergence of five focal populations (BS, DT, E8, PB, and X7) from a common ancestral source. The blue arrows represent estimated migration rates (gene flow) between specific population pairs, with values indicating the probability of migration per generation. The widths of the bars represent differences in effective population sizes (*N_e_*) estimated for each population. B) Results from a forward-time SLiM simulation tracking the pairwise *F_st_* between populations DT and E8 across generations. The solid blue line represents the mean *F_st_*, with the shaded area indicating the 95% confidence interval. The red dashed line denotes a differentiation threshold of 0.09.

A comparison between embryonic period differentiation (*Q_st_* EP) and neutral genetic differentiation (*F_st_*) shows a significant signature of divergent selection. The *Q_st_* for embryonic period is approximately 0.35, a value that is orders of magnitude higher than the mean overall *F_st_*of approximately 0.01 (Figure 5). Importantly, the 95% confidence intervals for these two metrics do not overlap, with the lower bound of *Q_st_* remaining well above the upper bound of *F_st_*.

**Figure 5:**
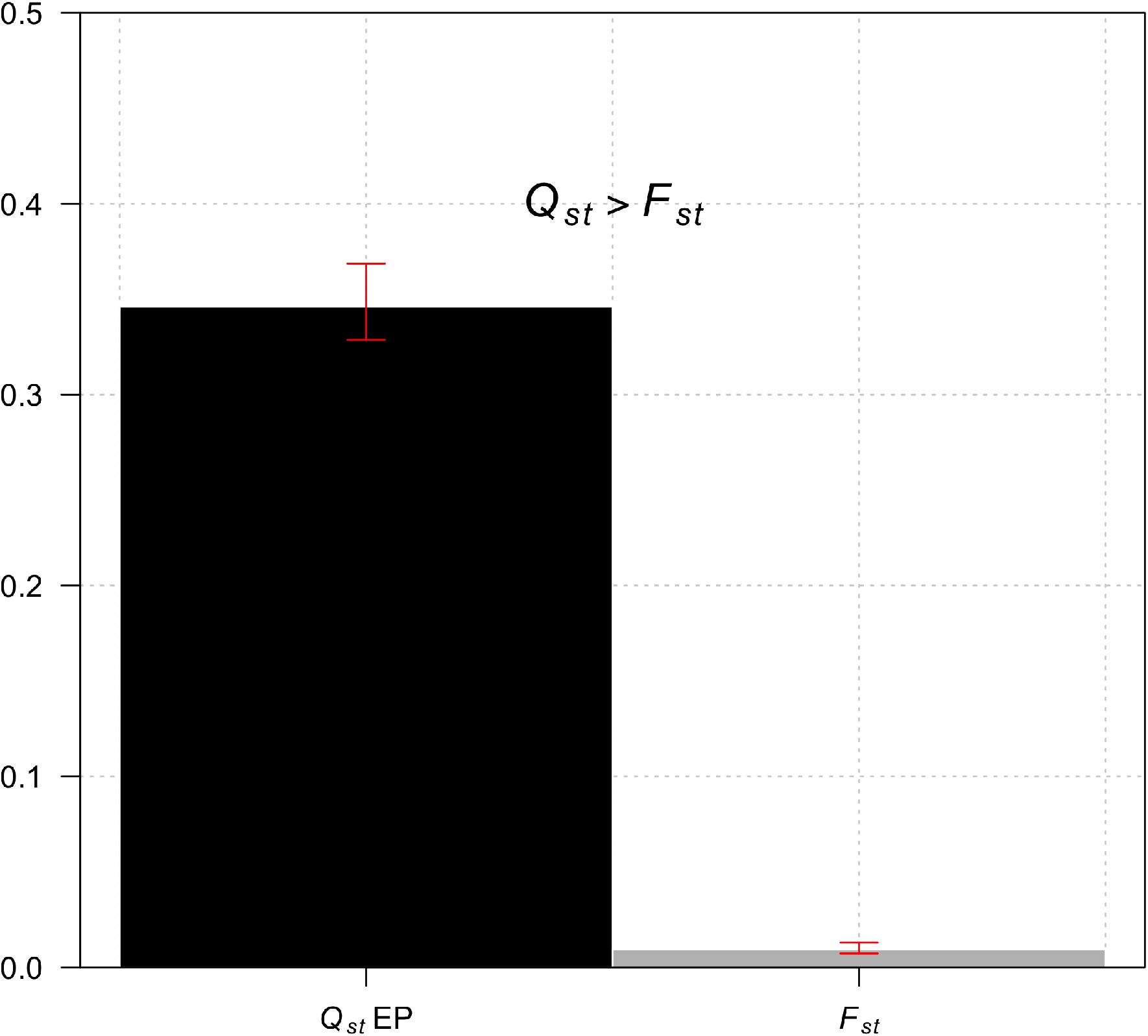
Comparison of phenotypic differentiation (*Q_st_*) and neutral genetic differentiation (*F_st_*) across the study populations. Red error bars represent the 95% confidence intervals around the mean value.

## Discussion

The genome annotation of the wood frog demonstrates high levels of completeness and contiguity, with BUSCO scores exceeding 90% (approximately 70% single-copy and 20% duplicated). These results compare favorably to other recent high-quality ranid assemblies, such as the chromosome-level assembly of the Common Frog (*Rana temporaria*), which achieved a BUSCO completeness of 95.2% for its annotated gene set (Streicher et al., 2021). Notably, the results of the wood frog annotation exhibits a significantly higher proportion of duplicated BUSCOs compared to the 2.4% duplication rate observed in the common frog and the 2.5% reported for the most recent American Bullfrog assembly (*Aquarana catesbeiana*; Zhang et al., 2025). This enrichment in duplicated orthologs likely reflects the complex genomic architecture of large-genome amphibians or lineage-specific gene family expansions. Furthermore, our results represent a drastic improvement over earlier draft assemblies in the lineage; for instance, the initial 2017 *A. catesbeiana* draft recovered only 45.3% complete BUSCOs (Hammond et al., 2017). The high completeness observed in our data, mirroring the 95.8% completeness seen in recent Neotropical Treefrog assemblies like *Dendropsophus ebraccatus* (NCBI RefSeq assembly GCF_027789765.1), underscores the robustness of our sequencing and annotation pipeline in capturing the intricate genomic landscape of the Ranidae family.

The availability of a fully annotated wood frog genome now enables us to accurately examine the species’ adaptive landscape that had previously remained uncharacterized. The detection of an outlier locus such as textitRab28 provides new evidence of regions potentially under divergent selection across the genome of the wood frog. The observation that the frequencies of these locus spikes at a single pond is intriguing and deserves further study since the pond has unusual combination of characteristics (low canopy cover and relatively large area). It is possible that this signature is associated with the previously documented countergradient variation in wood frogs influenced by canopy cover (Skelly, 2004). While previous studies in other species have focused on immune gene expression (e.g., MHC class *IIβ*) as a driver of local adaptation (Hernandez-Gomez et al., 2019), our findings suggest that structural genes like *Rab28* are equally vital. These genes likely work in tandem with metabolic adaptations such as the accumulation of cryoprotective glucose to allow wood frogs to persist in ecological conditions that would be lethal to other North American amphibians (Bay et al., 2018).

While the identification of specific outlier loci could be initially interpreted as a potential for local adaptation, their impact is most intuitively understood through their contribution to the fitness of individuals. Because we have no evidence of a clear link between these loci and a specific phenotype, we rather compared the differentiation in embryonic development (*Q_st_*) with neutral genetic differentiation (*F_st_*) to establish a robust framework for distinguishing between natural selection and genetic drift as drivers of phenotypic variation in the wood frog (Whitlock, 2008). We focused on embryonic development as it is considered a critical life-history trait that directly affects the fitness of the species (Skelly, 2004; Lind and Johansson, 2007; Le Sage et al., 2021). Quantifying *Q_st_* is also critical in this context because it may capture the cumulative effect of adaptive alleles responsible for the evolution of polygenic traits. Interestingly, some studies on amphibian populations have found that phenotypic traits often evolve in a neutral fashion (*Q_st_≈ F_st_*), suggesting that drift is the primary driver of variation in smaller, fragmented populations (Johansson et al., 2007; Leinonen et al., 2013). However, our results align more closely with research on wood frogs that identifies strong adaptive responses to environmental gradients, such as desiccation risk and temperature fluctuations (Lind and Johansson, 2007). Similar findings of *Q_st_ > F_st_* have been reported in life-history traits (Le Sage et al., 2021), suggesting that selection can overwhelm the effect of neutral forces among traits linked to the fitness of individuals. The evidence of this study strengthens the case that wood frog populations are fine-tuned to their local environment.

Although the effect of selection has been sufficiently strong to cause genomic and phenotypic divergence in the wood frog, our observation that populations maintain negligible genetic differentiation is intriguing. Previous studies suggest that wood frog populations can exhibit negligible genetic structure across heterogeneous landscapes, likely due to juvenile dispersal events (Crosby et al., 2009). By contrast, significant fine-scale population structure in wood frogs can be detected particularly in areas where agricultural development or high-traffic roads prevent connectivity (Newman and Squire, 2001). Our analysis indicates that the lack of broad-scale population structure likely results from a recent colonization event with ongoing gene flow among populations, limiting the time available for genetic drift to accumulate substantial differentiation. Supporting this idea, the SLiM simulation suggests that it might take approximately 400-500 generations for populations DT and E8 to reach a pairwise *F_st_* of 0.09. However, the time required for these populations to reach an *F_st_*of 0.05, a value that has been referred to as significant genetic differentiation for the species in a similar environment, is approximately 180 generations (Skibbe et al., 2021). Our results may reflect an incipient case of genomic islands of divergence in the wood frog, where selection maintains adaptive differences despite high levels of genetic connectivity. In a landscape characterized by ongoing gene flow, the majority of the genome is homogenized by the constant exchange of alleles through migration. However, localized regions (or “islands”) resist this homogenization due to the strength of divergent selection (Wolf and Ellegren, 2017). These islands typically harbor loci that are crucial for local adaptation, such as those governing physiological responses to environmental extremes or reproductive timing. When the selective advantage of an allele in a specific habitat outweighs the rate of migration, these regions remain highly differentiated and appear as significant outliers in genome-wide scans (e.g., Marques et al., 2017).

Overall, the fully annotated genome of the wood frog helps us better understand how specific genes can track ecological gradients, enabling us to provide insights into the adaptive ability of the species at a microgeographic scale. This genomic signature of selection together with a remarkable phenotypic differentiation strongly suggests that natural selection overcomes the power of genetic drift, even in a landscape characterized by a recent colonization and significant connectivity. Future research should aim to move from inferential models to the direct quantification of the evolutionary forces at play. A critical next step involves designing experiments that allow for the empirical measurement of selection coefficients (*s*) and migration rates (*m*) to precisely model the selection-migration balance across heterogeneous habitats. Given that the dispersal capacity and fine-scale movement patterns of wood frogs remain poorly understood, a long-term tracking experiment based on radio telemetry or high-resolution PIT tagging is necessary to validate our hypotheses regarding gene flow and its role in homogenizing neutral genomic regions. Coupling these ecological tracking data with experimental translocations would provide a more accurate test of the fitness advantages conferred by locally adapted alleles, ultimately offering a more granular understanding of how this species persists across its broad and often extreme environmental range.

## Acknowledgements

TBD

## Data Accessibility Statement

A fully reproducible workflow of the data analyses, including R scripts and additional supporting material, is available in the following repositories: Github https://dylan-padilla.github.io/YMF2018/. A dryad link will be available upon acceptance:.

## Supplementary material

**Figure S1:**
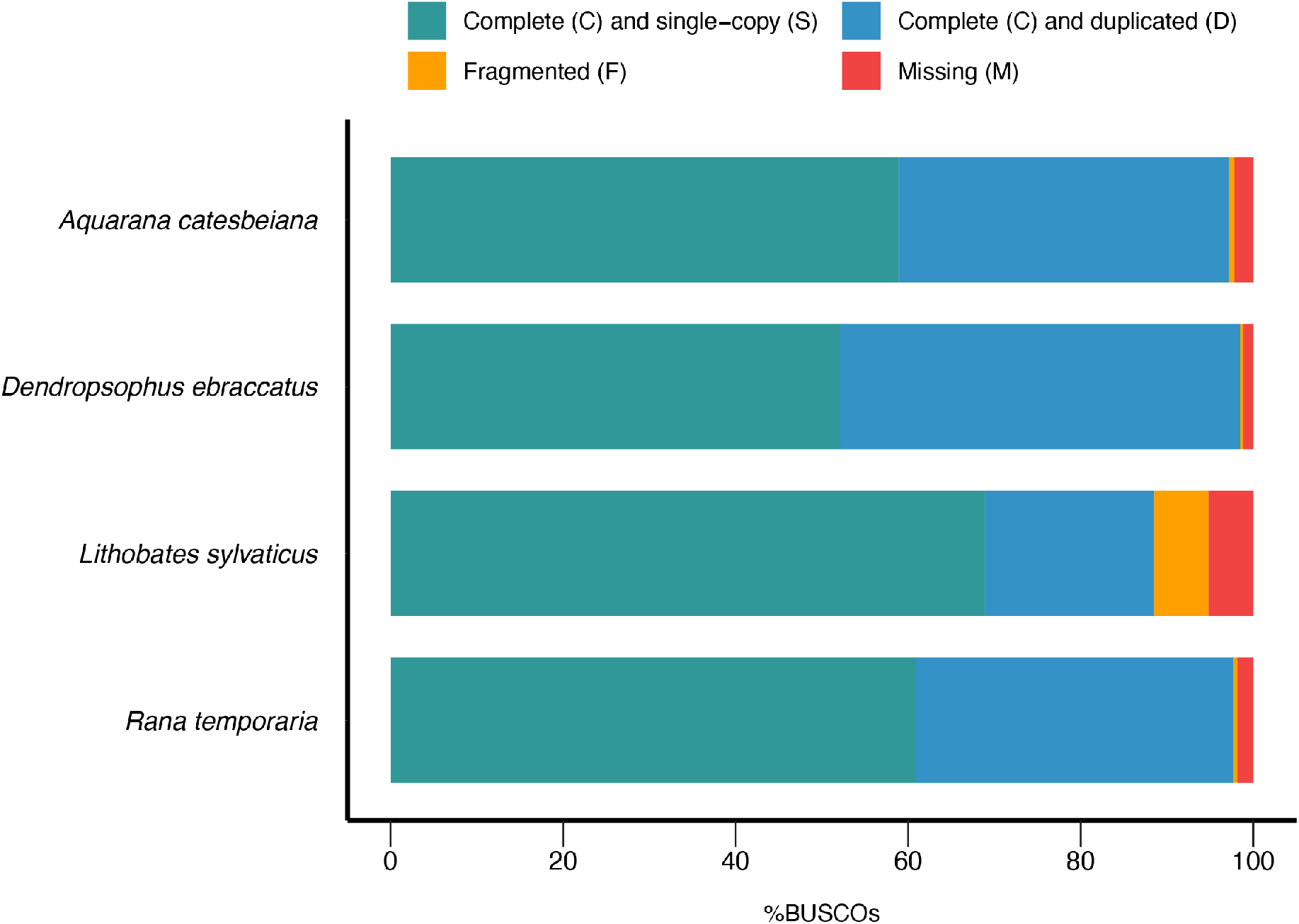
Comparative assessment of genome assembly and annotation completeness using Benchmarking Universal Single-Copy Orthologs (BUSCO). The bar chart illustrates the percentage of recovered conserved orthologs for *Lithobates sylvaticus* in relation to three reference anuran species: *Aquarana catesbeiana*, *Dendropsophus ebraccatus*, and *Rana temporaria*.

## References

Arietta, A. A. and Skelly, D. K. (2021). Rapid microgeographic evolution in response to climate change. Evolution, 75(11):2930–2943.

Bates, D., Mächler, M., Bolker, B., and Walker, S. (2015). Fitting linear mixed-effects models using lme4. Journal of Statistical Software, 67(1):1–48.

Bay, R. A., Harrigan, R. J., Underwood, V. L., Gibbs, H. L., Smith, T. B., and Ruegg, K. (2018). Genomic signals of selection predict climate-driven population declines in a migratory bird. Science, 359(6371):83–86.

Berven, K. A. and Grudzien, T. A. (1990). Dispersal in the wood frog (rana sylvatica): implications for genetic population structure. Evolution, 44(8):2047–2056.

Bruuna, T., Li, H., Guhlin, J., Honsel, D., Herbold, S., Stanke, M., Nenasheva, N., Ebel, M., Gabriel, L., and Hoff, K. J. (2023). Galba: genome annotation with miniprot and augustus. BMC bioinformatics, 24(1):327.

Buchfink, B., Xie, C., and Huson, D. H. (2015). Fast and sensitive protein alignment using diamond. Nature methods, 12(1):59–60.

Camacho, C., Coulouris, G., Avagyan, V., Ma, N., Papadopoulos, J., Bealer, K., and Madden, T. L. (2009). Blast+: architecture and applications. BMC bioinformatics, 10(1):421.

Clegg, S. M., Degnan, S. M., Kikkawa, J., Moritz, C., Estoup, A., and Owens, I. P. (2002). Genetic consequences of sequential founder events by an island-colonizing bird. Proceedings of the National Academy of Sciences, 99(12):8127–8132.

Coffman, A. J., Hsieh, P. H., Gravel, S., and Gutenkunst, R. N. (2016). Computationally efficient composite likelihood statistics for demographic inference. Molecular biology and evolution, 33(2):591–593.

Crawford, A. J. (2003). Relative rates of nucleotide substitution in frogs. Journal of Molecular Evolution, 57(6):636–641.

Crosby, M. K. A., Licht, L. E., and Fu, J. (2009). The effect of habitat fragmentation on finescale population structure of wood frogs (rana sylvatica). Conservation Genetics, 10(6):1707–1718.

Dainat, J., Hereñú, D., and Pucholt, P. (2020). Agat: Another gff analysis toolkit to handle annotations in any gtf. GFF format, 10.

Eaton, D. A. and Overcast, I. (2020). ipyrad: Interactive assembly and analysis of radseq datasets. Bioinformatics, 36(8):2592–2594.

Ewels, P., Magnusson, M., Lundin, S., and Käller, M. (2016). Multiqc: summarize analysis results for multiple tools and samples in a single report. Bioinformatics, 32(19):3047–3048.

Excoffier, L., Marchi, N., Marques, D. A., Matthey-Doret, R., Gouy, A., and Sousa, V. C. (2021). fastsimcoal2: demographic inference under complex evolutionary scenarios. Bioinformatics, 37(24):4882–4885.

Frichot, E. and François, O. (2015). Lea: An r package for landscape and ecological association studies. Methods in ecology and evolution, 6(8):925–929.

Frichot, E. and Francois, O. (2015). LEA: an R package for Landscape and Ecological Association studies. Methods in Ecology and Evolution.

Gruber, B., Unmack, P. J., Berry, O. F., and Georges, A. (2018). dartr: An r package to facilitate analysis of snp data generated from reduced representation genome sequencing. Molecular ecology resources, 18(3):691–699.

Gutenkunst, R. N., Hernandez, R. D., Williamson, S. H., and Bustamante, C. D. (2009). Inferring the joint demographic history of multiple populations from multidimensional snp frequency data. PLoS genetics, 5(10):e1000695.

Haller, B. C., Ralph, P. L., and Messer, P. W. (2026). Slim 5: Eco-evolutionary simulations across multiple chromosomes and full genomes. Molecular Biology and Evolution, 43(1):msaf313.

Hammond, S. A., Warren, R. L., Vandervalk, B. P., Kucuk, E., Khan, H., Gibb, E. A., Pandoh, P., Kirk, H., Zhao, Y., Jones, M., et al. (2017). The north american bullfrog draft genome provides insight into hormonal regulation of long noncoding rna. Nature communications, 8(1):1433.

Hernandez-Gomez, O., Kimble, S. J., Hua, J., Wuerthner, V. P., Jones, D. K., Mattes, B. M., Cothran, R. D., Relyea, R. A., Meindl, G. A., and Hoverman, J. T. (2019). Local adaptation of the mhc class ii*β* gene in populations of wood frogs (lithobates sylvaticus) correlates with proximity to agriculture. Infection, Genetics and Evolution, 73:197–204.

Hoff, K. J. and Stanke, M. (2019). Predicting genes in single genomes with augustus. Current protocols in bioinformatics, 65(1):e57.

Johansson, M., Primmer, C. R., and Merilä, J. (2007). Does habitat fragmentation reduce fitness and adaptability? a case study of the common frog (rana temporaria). Molecular Ecology, 16(13):2693–2700.

Jumentier, B., Caye, K., Heude, B., Lepeule, J., and François, O. (2022). Sparse latent factor regression models for genome-wide and epigenome-wide association studies. Statistical Applications in Genetics and Molecular Biology, 21(1):20210035.

Junge, C., Vøllestad, L., Barson, N., Haugen, T., Otero, J., Sætre, G., Leder, E., and Primmer, C. (2011). Strong gene flow and lack of stable population structure in the face of rapid adaptation to local temperature in a spring-spawning salmonid, the european grayling (thymallus thymallus). Heredity, 106(3):460–471.

Kamvar, Z. N., Tabima, J. F., and Grünwald, N. J. (2014). Poppr: an r package for genetic analysis of populations with clonal, partially clonal, and/or sexual reproduction. PeerJ, 2:e281.

Kealoha Freidenburg, L. and Skelly, D. K. (2004). Microgeographical variation in thermal preference by an amphibian. Ecology Letters, 7(5):369–373.

Le Sage, E. H., Duncan, S. I., Seaborn, T., Cundiff, J., Rissler, L. J., and Crespi, E. J. (2021). Ecological adaptation drives wood frog population divergence in life history traits. Heredity, 126(5):790–804.

Leinonen, T., Cano, J. M., Mäkinen, H., and Merilä, J. (2006). Contrasting patterns of body shape and neutral genetic divergence in marine and lake populations of threespine sticklebacks. Journal of evolutionary biology, 19(6):1803–1812.

Leinonen, T., McCairns, R. S., O’hara, R. B., and Merilä, J. (2013). Q st–f st comparisons: evolutionary and ecological insights from genomic heterogeneity. Nature Reviews Genetics, 14(3):179–190.

Lenormand, T. (2002). Gene flow and the limits to natural selection. Trends in ecology & evolution, 17(4):183–189.

Li, H. (2023). Protein-to-genome alignment with miniprot. Bioinformatics, 39(1):btad014.

Lind, M. I. and Johansson, F. (2007). The degree of adaptive phenotypic plasticity is correlated with the spatial environmental heterogeneity experienced by island populations of rana temporaria. Journal of evolutionary biology, 20(4):1288–1297.

Marques, D. A., Lucek, K., Haesler, M. P., Feller, A. F., Meier, J. I., Wagner, C. E., Excoffier, L., and Seehausen, O. (2017). Genomic landscape of early ecological speciation initiated by selection on nuptial colour. Molecular ecology, 26(1):7–24.

Merilä, J. and Crnokrak, P. (2001). Comparison of genetic differentiation at marker loci and quantitative traits. Journal of Evolutionary Biology, 14(6):892–903.

Mila, B., Smith, T. B., and Wayne, R. K. (2007). Speciation and rapid phenotypic differentiation in the yellow-rumped warbler dendroica coronata complex. Molecular Ecology, 16(1):159–173.

Newman, R. A. and Squire, T. (2001). Microsatellite variation and fine-scale population structure in the wood frog (rana sylvatica). Molecular Ecology, 10(5):1087–1100.

Nielsen, R. (2005). Molecular signatures of natural selection. Annu. Rev. Genet., 39(1):197–218.

Nosil, P., Funk, D. J., and Ortiz-Barrientos, D. (2009). Divergent selection and heterogeneous genomic divergence. Molecular ecology, 18(3):375–402.

Palo, J., O’hara, R. B., Laugen, A. T., Laurila, A., Primmer, C. R., and Merilä, J. (2003). Latitudinal divergence of common frog (rana temporaria) life history traits by natural selection: evidence from a comparison of molecular and quantitative genetic data. Molecular ecology, 12(7):1963–1978.

Paradis, E. (2010). pegas: an r package for population genetics with an integrated– modular approach. Bioinformatics, 26(3):419–420.

Purcell, S., Neale, B., Todd-Brown, K., Thomas, L., Ferreira, M. A., Bender, D., Maller, J., Sklar, P., De Bakker, P. I., Daly, M. J., et al. (2007). Plink: a tool set for whole-genome association and population-based linkage analyses. The American journal of human genetics, 81(3):559–575.

Richardson, J. L., Urban, M. C., Bolnick, D. I., and Skelly, D. K. (2014). Microgeographic adaptation and the spatial scale of evolution. Trends in ecology & evolution, 29(3):165–176.

Savolainen, O., Lascoux, M., and Merilä, J. (2013). Ecological genomics of local adaptation. Nature Reviews Genetics, 14(11):807–820.

Skelly, D. and Freidenburg, L. (2000). Effects of beaver on the thermal biology of an amphibian. Ecology Letters, 3(6):483–486.

Skelly, D. K. (2004). Microgeographic countergradient variation in the wood frog, rana sylvatica. Evolution, 58(1):160–165.

Skibbe, J. R., Farrar, J., Watson, K., and Richter, S. C. (2021). Population genetics of wood frogs (lithobates sylvaticus) in an altered forested ridgetop wetland ecosystem in appalachia. Herpetological Conservation and Biology, 16(1):1–10.

Stanke, M., Schöffmann, O., Morgenstern, B., and Waack, S. (2006). Gene prediction in eukaryotes with a generalized hidden markov model that uses hints from external sources. BMC bioinformatics, 7(1):62.

Storey, J. D. and Tibshirani, R. (2003). Statistical significance for genomewide studies. Proceedings of the National Academy of Sciences, 100(16):9440–9445.

Streicher, J. W., of Life, W. S. I. T., of Life Consortium, D. T., et al. (2021). The genome sequence of the common frog, rana temporaria linnaeus 1758. Wellcome Open Research, 6:286.

Sun, Y.-B., Xiong, Z.-J., Xiang, X.-Y., Liu, S.-P., Zhou, W.-W., Tu, X.-L., Zhong, L., Wang, L., Wu, D.-D., Zhang, B.-L., et al. (2015). Whole-genome sequence of the tibetan frog nanorana parkeri and the comparative evolution of tetrapod genomes. Proceedings of the National Academy of Sciences, 112(11):E1257–E1262.

Tate, R., Hall, B., DeRego, T., and Geib, S. (2014). Annie: The annotation information extractor (version 1.0).

Weir, B. S. and Cockerham, C. C. (1984). Estimating f-statistics for the analysis of population structure. evolution, pages 1358–1370.

Whitlock, M. C. (2008). Evolutionary inference from qst. Molecular ecology, 17(8):1885–1896.

Wolf, J. B. and Ellegren, H. (2017). Making sense of genomic islands of differentiation in light of speciation. Nature Reviews Genetics, 18(2):87–100.

Wright, S. (1931). Evolution in mendelian populations. Genetics, 16(2):97.

Zhang, K., Zhang, Y., Tian, Y., Xu, B., Jiang, X., Qin, Z., Liu, C., and Lin, L. (2025). A chromosome-level genome assembly of the american bullfrog (aquarana catesbeiana). Scientific Data, 12(1):413.

